# Soil-based disease bioassay for the study of rhizogenic *Agrobacterium*-tomato interactions

**DOI:** 10.64898/2026.04.14.718410

**Authors:** Savio D. Rodrigues, Nuri Kim, Joram Moons, Hans Rediers, Laurens Pauwels, Barbara De Coninck

## Abstract

Hairy root disease (HRD), caused by rhizogenic *Agrobacterium*, is an economically important disease affecting hydroponic tomato (*Solanum lycopersicum* L.) production worldwide. HRD-affected plants show extensive root proliferation, resulting in decreased energy expenditure towards fruit production. Host plant susceptibility to rhizogenic *Agrobacterium* is typically evaluated through artificial wounding-based infection bioassays. However, under natural infection settings, rhizogenic *Agrobacterium* can induce disease symptoms without deliberate, artificial wounding. We developed a soil-based, non-wounding bioassay that closely mimics natural rhizosphere interactions and permits quantitative and qualitative assessment of HRD symptoms. The assay measured root dry weight, documented agravitropic root development typical of HRD and confirmed *in planta* T-DNA gene expression using reverse transcriptase quantitative polymerase chain reaction (RT-qPCR). We used this bioassay to evaluate disease symptoms towards rhizogenic *Agrobacterium* in tomato cv. ‘Moneymaker’ and the rootstocks ‘Optifort’, ‘Maxifort’, and ‘Arnold’. ‘Optifort’ and ‘Maxifort’ exhibited significantly higher root biomass than ‘Arnold’ and ‘Moneymaker’, indicating more pronounced symptom development. The bioassay also differentiated virulence levels amongst various rhizogenic *Agrobacterium* strains isolated from HRD-affected plants. Together, these results show that our soil-based bioassay provides a robust and ecologically relevant platform for screening tomato genotypes and comparing virulence levels of rhizogenic *Agrobacterium* strains supporting resistance breeding and disease management efforts.

## 4. Introduction

High-tech greenhouses in temperate regions enable precise control of climate and plant nutrition, supporting intensive production of economically important crops (Nemali 2022; Thomas et al. 2024; Wittwer and Castilla 1995). Within these greenhouses, crops such as tomato (*Solanum lycopersicum* L.), bell pepper (*Capsicum annuum* L.), cucumber (*Cucumis sativus* L.), eggplant (*Solanum melongena* L.), and melon (*Cucumis melo* L.) are often cultivated using substrate-based hydroponic set-ups. This supports intensive cultivation practices by optimally balancing nutrient delivery and root aeration, ensuring high fruit yields and efficient resource use (Chatterjee et al. 2025; Jensen 1997; Tuxun et al. 2025). Although hydroponic set-ups limit many important soil-borne pests and pathogens (Rajendran et al. 2024), pathogens such as *Pythium, Fusarium, Verticillium, Phytophthora*, and *Rhizoctonia* continue to cause significant losses in commercial production (Bardin and Gullino 2020; Fussy and Papenbrock 2022; Os and Benoit 1999; Paulitz and Bélanger 2001; Vlasselaer et al. 2024). To improve crop performance, growers commonly graft scions onto interspecific hybrid rootstocks. This provides vigor, abiotic stress tolerance, and/or disease resistance to the grafted scion (Alqardaeai et al. 2025; Coşkun 2023; Djidonou et al. 2013; King et al. 2010; Mozafarian et al. 2020; Spanò et al. 2020). For tomato plants, commercially available rootstocks such as ‘Maxifort’, ‘Beaufort’, ‘Optifort’, ‘DRO141TX’, ‘Fortamino’, and ‘Arnold’ combine *S. lycopersicum* and *Solanum habrochaites* genetics (Cortada et al. 2008; Ludeking and Janse 2011; Vanlommel et al. 2019, 2020).

Since the late 1970’s, hairy root disease (HRD) (or colloquially known as root mat disease and crazy root disease), has been observed to globally impact both, soil and hydroponically-grown Solanaceae (tomatoes, bell peppers, and egg plants) and Cucurbitaceae (cucumber and melon) crops (Bosmans et al. 2015, 2017b; Eberle et al. 2020; Han et al. 2021; Ignatov et al. 2016; Janse 2008; Sawada and Azegami 2014; Shiomi et al. 1987; Vanlommel et al. 2020; Vargas et al. 2020; Warabieda et al. 2021; Weller et al. 2000b, 2000a). In Flanders, 45% of commercial tomato greenhouses were reported to be affected by the disease (Vanlommel et al. 2020). HRD is attributed to the gram-negative, *Agrobacterium* spp. harboring a root-inducing (Ri) plasmid (hereafter, referred to as rhizogenic *Agrobacterium*). During infection, rhizogenic *Agrobacterium* transfers and integrates a part of its Ri plasmid (known as transfer DNA or T-DNA) into the host’s genome, resulting in irreversible disease symptoms. Plants with HRD show excessive agravitropic root proliferation, altered shoot growth, reduced fruit size, and yield losses of up to 15% in some plant varieties (Bosmans et al. 2017b; Weller et al. 2000a). Excessive root proliferation further impairs irrigation and can promote occurrence of secondary infections such as *Pythium*-induced root rot (Bosmans et al. 2017b; de Freitas and Taylor 2023; Weller et al. 2000b). Because HRD results from genomic integration of the Ri T-DNA, no curative treatments exist. Current management measures include *a)* strict phytosanitary practices (physical and/or chemical disinfection of irrigation systems using UV-C, heat, chlorine-based disinfectants, or hydrogen peroxide); and *b)* modified cultivation strategies (choice of rootstock; substrate type; irrigation regimes) (Bosmans et al. 2017a, 2017b; Bourigault et al. 2021; Fortuna et al. 2023; de Freitas and Taylor 2023; Vanlommel et al. 2020; Vargas et al. 2021).

Several functional bioassays have been developed to study rhizogenic *Agrobacterium* and assess host susceptibility or *Agrobacterium* virulence (Christey 1997, 2001; Cleene and De Ley 1981; Collier et al. 2005; Desmet et al. 2019; Hansen et al. 1989; Liao 1978; Porter and Flores 1991; White et al. 1985). Most assays rely on deliberate, artificial wounding of plant tissue prior to rhizogenic *Agrobacterium* inoculation as wounding facilitates plant transformation (Christey and Braun 2004). This wounding-based strategy, however, has limited ecological relevance since natural greenhouse infections occur without artificial wounding. A few non-wounding dependent bioassays using soil or substrate-grown plants have also been reported (de Freitas and Taylor 2023; Kawaguchi et al. 2008; Ludeking and Janse 2011; Vanlommel et al. 2020; Vargas et al. 2020). These assays compared qualitative parameters such as disease incidence (measured as extent of development of agravitropic growing roots or presence/absence of hairy roots) (de Freitas and Taylor 2023; Vanlommel et al. 2020; Vargas et al. 2020) but did not quantify biomass development, which is an important HRD-relevant agronomic trait linked to crop production. Importantly, although qualitative observations suggest that different tomato rootstocks vary in HRD susceptibility (Ludeking and Janse 2011; Vanlommel et al. 2019), no quantitative comparisons have been reported. Additionally, despite known strain-to-strain variation in the ability of rhizogenic *Agrobacterium* to induce hairy roots (Desmet et al. 2019), their relative virulence has not been assessed on tomato or its rootstocks under natural infection settings.

Here, we developed a non-wounding, soil-based disease bioassay to characterize tomato HRD under natural infection settings. The assay supports quantitative and qualitative evaluation of HRD symptoms. We used the bioassay to compare *a)* susceptibilities of tomato cv. ‘Moneymaker’, and three tomato rootstocks, ‘Optifort’, ‘Maxifort’, and ‘Arnold’, to HRD; and *b)* virulence levels of a collection of five rhizogenic *Agrobacterium* strains isolated from HRD-affected plant roots on ‘Optifort’.

## 5. Materials and Methods

### 5.1 Bacterial strains, media, and conditions

All rhizogenic *Agrobacterium* strains used in this study are listed in Table 1. Briefly, rhizogenic *Agrobacterium* strains NCPPB2659 [hereafter, referred to as K599; belongs to *Agrobacterium* spp. genomospecies group G21; (Singh et al. 2021)], NCPPB4062 [hereafter, referred to as 4062; belongs to *Agrobacterium deltaense* genomospecies group G7 (Bosmans et al. 2015)], ST15.13/012 [hereafter referred to as 012; belongs to *Agrobacterium salinitolerans* genomospecies group G9 (Bosmans et al. 2015)], ST15.13/097 [hereafter, referred to as 097; belongs to *Agrobacterium salinitolerans* genomospecies group G9 (Vargas et al. 2024)], ST15.13/057 [hereafter, referred to as 057; belongs to *Agrobacterium* genomospecies group G25 (Vargas et al. 2024)], and the disarmed rhizogenic *Agrobacterium* strain, 18r12v-Δ*recA*::*tetA* [hereafter referred to as 18r12v; disarmed variant of K599 lacking T-DNA; *recA* was replaced by tetracycline resistance gene (Collier et al. 2016)] were used in this study. These strains were grown from glycerol stocks on solid Yeast Extract Peptone (YEP) medium (10 g/L Yeast Extract, 10 g/L Peptone, 5 g/L NaCl, 15 g/L Agar) at 28 °C. Single colonies were inoculated into 20 mL liquid YEP medium and grown for 24 h at 28 °C. These served as seed cultures for inoculating 500 mL liquid YEP medium in baffled flasks and were incubated for 24 h at 28 °C. For all rhizogenic *Agrobacterium* inoculations (i.e., soil drenching/re-inoculation), bacterial cells were harvested by centrifugation at 4000 x g for 10 min and resuspended in ½ strength Murashige and Skoog (MS) liquid medium (2.151 g/L MS basal salts, 0.5 g/L 2-(N-morpholino)ethanesulfonic acid (MES) monohydrate, pH 5.6 - 5.8) to a final O.D._600_ of 0.1 (≈ 1 * 10^8^ colony forming units/mL [CFU/mL]).

**Table 1.**
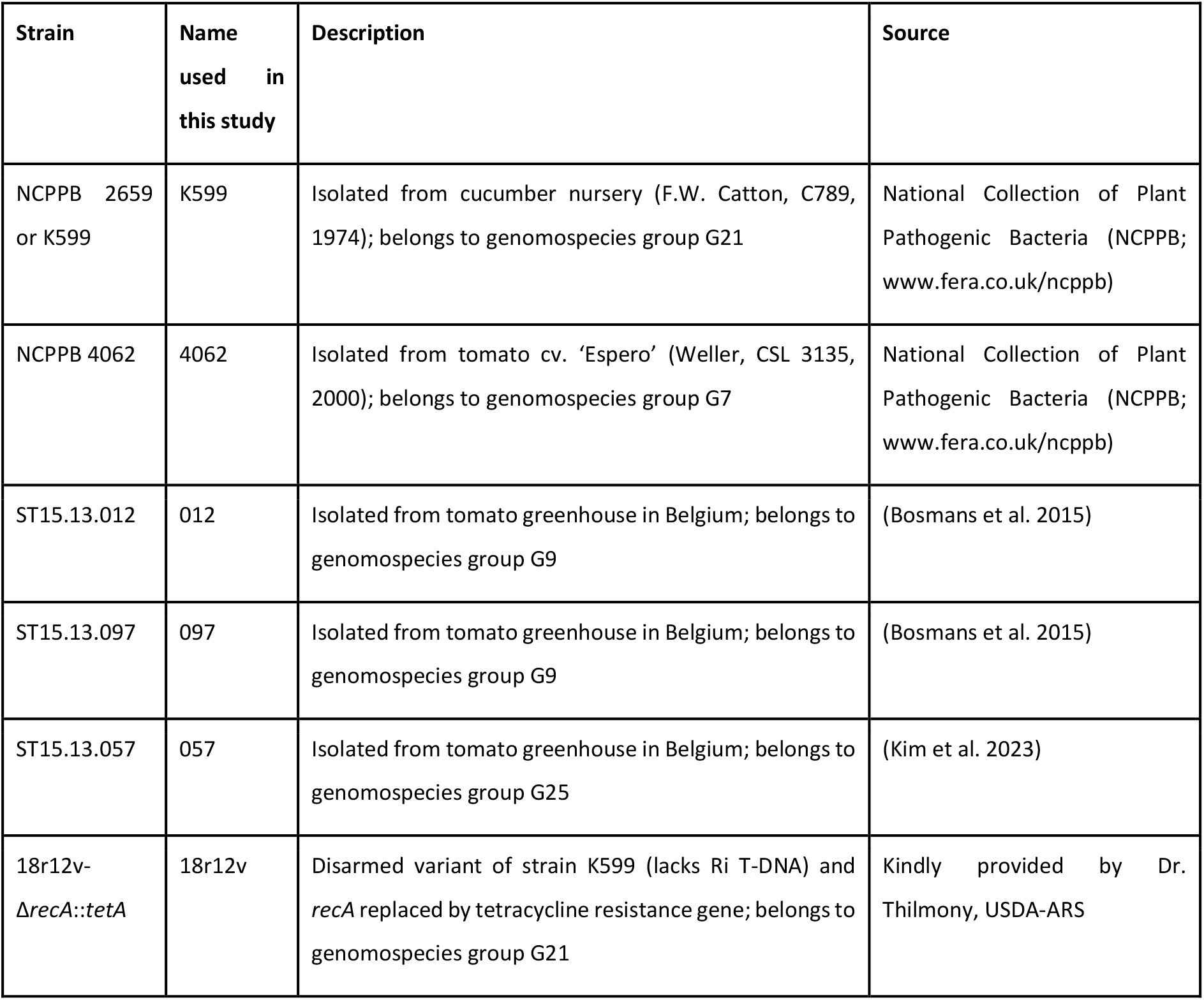
Bacterial strains used in this study.

| Strain | Name used in this study | Description | Source |
| --- | --- | --- | --- |
| NCPPB 2659 or K599 | K599 | Isolated from cucumber nursery (F.W. Catton, C789, 1974); belongs to genomospecies group G21 | National Collection of Plant Pathogenic Bacteria (NCPPB; www.fera.co.uk/ncppb) |
| NCPPB 4062 | 4062 | Isolated from tomato cv. 'Espero' (Weller, CSL 3135, 2000); belongs to genomospecies group G7 | National Collection of Plant Pathogenic Bacteria (NCPPB; www.fera.co.uk/ncppb) |
| ST15.13.012 | 012 | Isolated from tomato greenhouse in Belgium; belongs to genomospecies group G9 | (Bosmans et al. 2015) |
| ST15.13.097 | 097 | Isolated from tomato greenhouse in Belgium; belongs to genomospecies group G9 | (Bosmans et al. 2015) |
| ST15.13.057 | 057 | Isolated from tomato greenhouse in Belgium; belongs to genomospecies group G25 | (Kim et al. 2023) |
| 18r12v-*ΔrecA::tetA* | 18r12v | Disarmed variant of strain K599 (lacks Ri T-DNA) and *recA* replaced by tetracycline resistance gene; belongs to genomospecies group G21 | Kindly provided by Dr. Thilmony, USDA-ARS |

### 5.2 Plant varieties and growth conditions

Tomato (*S. lycopersicum* L.) cv. ‘Moneymaker’ (hereafter, referred to as ‘Moneymaker’; Garden Seeds, The Netherlands), and tomato rootstocks, ‘Maxifort’ (*S. lycopersicum* x *S. habrochaites*; De Ruiter Seeds, The Netherlands), ‘Optifort’ (*S. lycopersicum* x *S. habrochaites*; De Ruiter Seeds, The Netherlands), and ‘Arnold’ (*S. lycopersicum* x *S. habrochaites*; Syngenta Seeds, The Netherlands) were used in this study. For the rootstocks, primed, non-treated seeds were employed. Seeds were surface sterilized prior to germination as described in Ron et al. (2014) with some modifications. Briefly, seeds were placed in 70% ethanol for 10 min followed by washing with sterile MilliQ water. The seeds were then soaked in 50% commercial bleach (10° concentration; diluted in MilliQ water) for 15 min and washed five times with sterile MilliQ water. The seeds were transferred aseptically over moist filter papers (Whatmann qualitative filter paper grade 1) in Petri dishes, sealed with micropore tape and placed in the dark at 25 °C. After four days, the Petri dishes were transferred to a growth chamber (25 °C; 16-8 photoperiod [16 h light, 8 h dark]) until transplanting.

Seedlings were transplanted into a soil-sand mixture drenched with rhizogenic *Agrobacterium* and grown for nine to twelve weeks at 25 °C with 70% relative humidity and 120 µmol.m^-2^.s^-1^ of LED light (Sunritek VS10 FSM). Plants received weekly bottom watering with modified Hoagland’s solution (0.5 g/L MgSO_4_.7H_2_O, 0.27 g/L KH_2_PO_4_, 0.2 g/L KNO_3_, 0.1 g/L K_2_SO_4_, 0.5 g/L Ca(NO_3_)_2_.4H_2_O, 0.025 g/L FeEDTA Sodium salt, 0.0041 g/L H_3_BO_3_, 0.0037 g/L MnSO_4_.H_2_O, 0.0002 g/L CuCl_2_.2H_2_O, 0.0000825 g/L (NH_4_)_6_Mo_7_O_24_.4H_2_O, 0.000649 g/L ZnSO_4_.7H_2_O, pH 5.6-5.8). Additional weekly requirements for irrigation were met with demineralized water.

### 5.3 Soil-based disease bioassay

The disease bioassay used in this study was adapted from Kawaguchi et al. (2008). A soil-sand mix was prepared using two parts sieved potting-soil (DCM zaai en stek potgrond terreau; DCM, The Netherlands) and one part sand (Rijnzand; Paesen, Belgium) (2:1 V/V). The mixture was drenched with rhizogenic *Agrobacterium* suspension to achieve 5 * 10^7^ CFU/g soil. Mock controls received the same volume of sterile ½ strength MS liquid medium. Approximately, 800-900 g of mock or rhizogenic *Agrobacterium*-drenched soil was filled in 1.5 L pots and seven-day-old seedlings were transplanted individually. Pots were covered with cling film for five days to maintain high relative humidity. Seven days after transplanting, plants were re-inoculated with an additional 5 * 10^7^ CFU/g by pouring inoculum directly over the soil surface. Mock controls received the same volume of sterile ½ strength MS liquid medium. Aluminum foil was used to cover the pot surface to prevent light exposure to developing roots. Plants were maintained for eight to eleven weeks after transplanting.

At the experimental endpoint, shoots were excised, and top surface of the potted soil was photographed. Shoot length, number of leaf nodes, and stem thickness were recorded. Roots were carefully harvested to minimize root breakage and washed to remove the soil-sand mix. The length of the root system was measured. About five roots (2-3 cm in length) were separately retained for RNA extractions. The remainder of the harvested roots were dried at 70 °C for seven days before recording dry weights (Fig. 1).

**Fig. 1.**
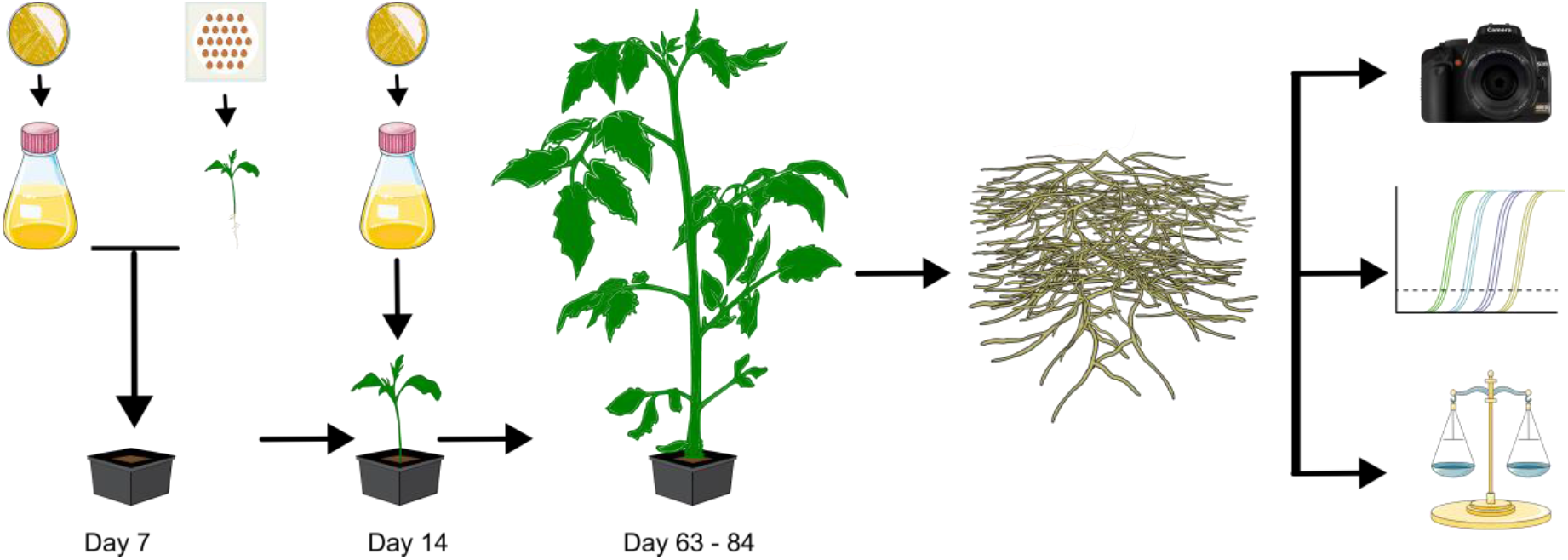
A non-wounding, soil-based infection assay to evaluate susceptibility of plant varieties and virulence levels of rhizogenic *Agrobacterium*. Sterile seven day-old tomato cv. ‘Moneymaker’ or interspecific hybrid rootstocks, ‘Arnold’, ‘Maxifort’, or ‘Optifort’, were transplanted to rhizogenic *Agrobacterium* drenched soil. 14-day-old seedlings were re-inoculated with respective rhizogenic *Agrobacterium* and symptom development was monitored in plants after nine to twelve weeks. At the end of the assay, the soil surface of individual pots was photographed followed by careful harvesting and cleaning of roots. Quantitative reverse-transcriptase polymerase chain reaction for rhizogenic *Agrobacterium* T-DNA genes and tomato housekeeping genes was performed on RNA extracted from ∼ five harvested roots (2-3 cm in length). The remainder of the harvested roots were dried in an oven at 70 °C for seven days and dry weights were measured. Illustration was constructed in Inkscape. (Adapted Petri dish with bacteria from petri-dish-with-bacteria-brown, culture flask from bath_flask, and weighing scale from scale-balanced by Servier https://smart.servier.com/ is licensed under CC-BY 3.0 Unported; Adapted plant pot from Brachypodium_Flowering_plant by Frédéric Bouché is licensed under CC-BY 4.0 Unported; Adapted seeds from seed_beetle_egg by DBCLS is licensed under CC-BY 4.0 Unported; Tomato seedling by Ciera Martinez is licensed under CC-BY 4.0 Unported; Adult tomato plants by EBI Gene Expression Group is licensed under CC-BY 4.0; Adapted root system by James-Lloyd is licensed under CC0; Camera by OpenClipart is licensed under CC0; qPCR_plot by Marcel Tisch is licensed under CC0)

### 5.4 Reverse-transcriptase quantitative PCR

Five individual roots (approximately 2-3 cm in length) were collected from every plant in separate tubes, flash-frozen in liquid nitrogen and stored at −80 °C until RNA was extracted. Total RNA was extracted from the roots using the Reliaprep™ RNA Tissue miniprep system kit (Promega) according to the manufacturer’s instructions for non-fibrous tissues. For complementary DNA (cDNA) synthesis, 500 ng DNase I-treated RNA was reverse transcribed using oligo(dT) primers (IDT, Belgium) and SuperScript™ IV Reverse Transcriptase (Thermo Fisher Scientific). cDNA was diluted threefold and used as template for reverse-transcriptase quantitative PCR (RT-qPCR) with SybrGreen™ (Invitrogen) on a CFX96 Touch Real-Time PCR Detection System (BioRad) (40 cycles of denaturation 95 °C for 15 sec, annealing 58.4 °C for 10 sec, and extension 72 °C for 1 min, with final melt curve analyses from 65 °C to 95 °C with increment of 0.5 °C for 5 sec and plate read). Expression of the reference gene *CLATHRIN ADAPTOR COMPLEXES MEDIUM SUBUNIT* (Sl*CAC*, Solyc08g006960) (Expósito-Rodríguez et al. 2008) confirmed RNA quality while the Ri T-DNA encoded *root oncogenic locus C* (*rolC*) was used to confirm the *in planta* expression of T-DNA genes. The primer sequences used for RT-qPCR are listed in Table 2. After RT-qPCR, amplicons were visualized by agarose gel electrophoresis to confirm amplification of the target genes.

**Table 2.**
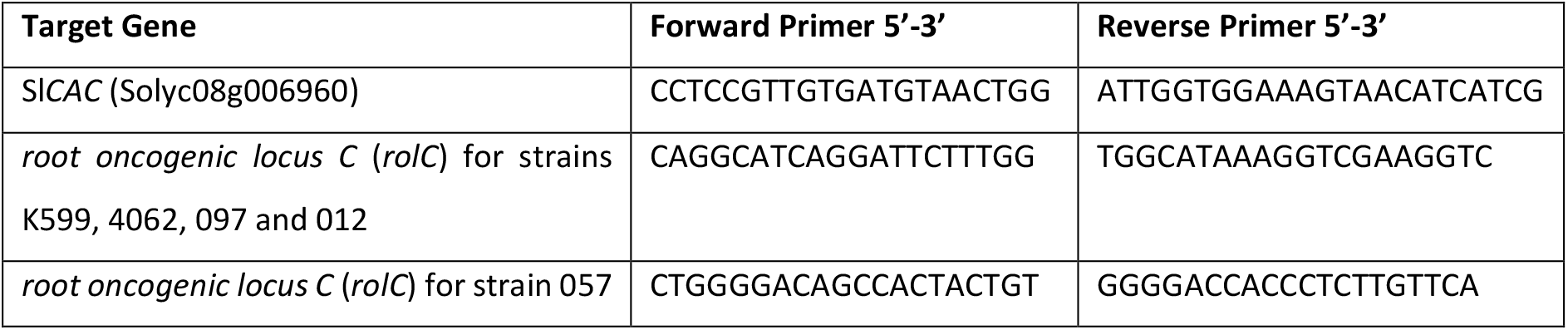
Primers utilized in this study.

### 5.5 Statistical analyses

All statistical analyses were performed in Graphpad Prism 8 and 10 for Windows. The normal distribution of datasets was evaluated using D’Agostino and Pearson, Anderson-Darling, and Shapiro-Wilk tests. Datasets showing lognormal distributions were transformed to log_10_ scales, and the distribution of the datasets was re-evaluated. Outliers identified from histograms of log_10_ transformed values were removed before analyses. Equality of variance were assessed using F test (for t-test) or Brown-Forsythe test and Bartlett’s test (for Ordinary One-Way ANOVA).For normally distributed transformed data with equal variance, Ordinary one-way ANOVA with Tukey’s *post hoc* test for multiple comparisons. For normally distributed transformed data with unequal variance, we used unpaired t-test with Welch’s correction with two-tailed *p*-value.

## 6 Results

### 6.1 ‘Optifort’ and ‘Maxifort’ show pronounced HRD symptoms following inoculation with rhizogenic *Agrobacterium*

We first performed a pilot soil-based disease bioassay using ‘Maxifort’ and rhizogenic *Agrobacterium* strain 057, to determine whether the assay captured quantifiable HRD-related plant parameters. ‘Maxifort’ and strain 057 were chosen for the pilot study as ‘Maxifort’ was reported to be susceptible to tomato HRD, while strain 057 has been reported to be highly virulent across different plant species (Rüter et al. 2024; Vanlommel et al. 2020; Vargas et al. 2020). At the assay endpoint (eight weeks post first inoculation), we measured shoot length, number of leaf nodes, stem thickness, root length, and root dry weight. Among these parameters, only root dry weight showed a significant 1.8-fold increase in strain 057-treated plants compared to the mock-treated plants (Fig 2A and *SI Appendix*, Fig. S1). This confirmed that root dry weight served as the most informative quantitative marker for HRD under our experimental conditions.

**Fig. 2.**
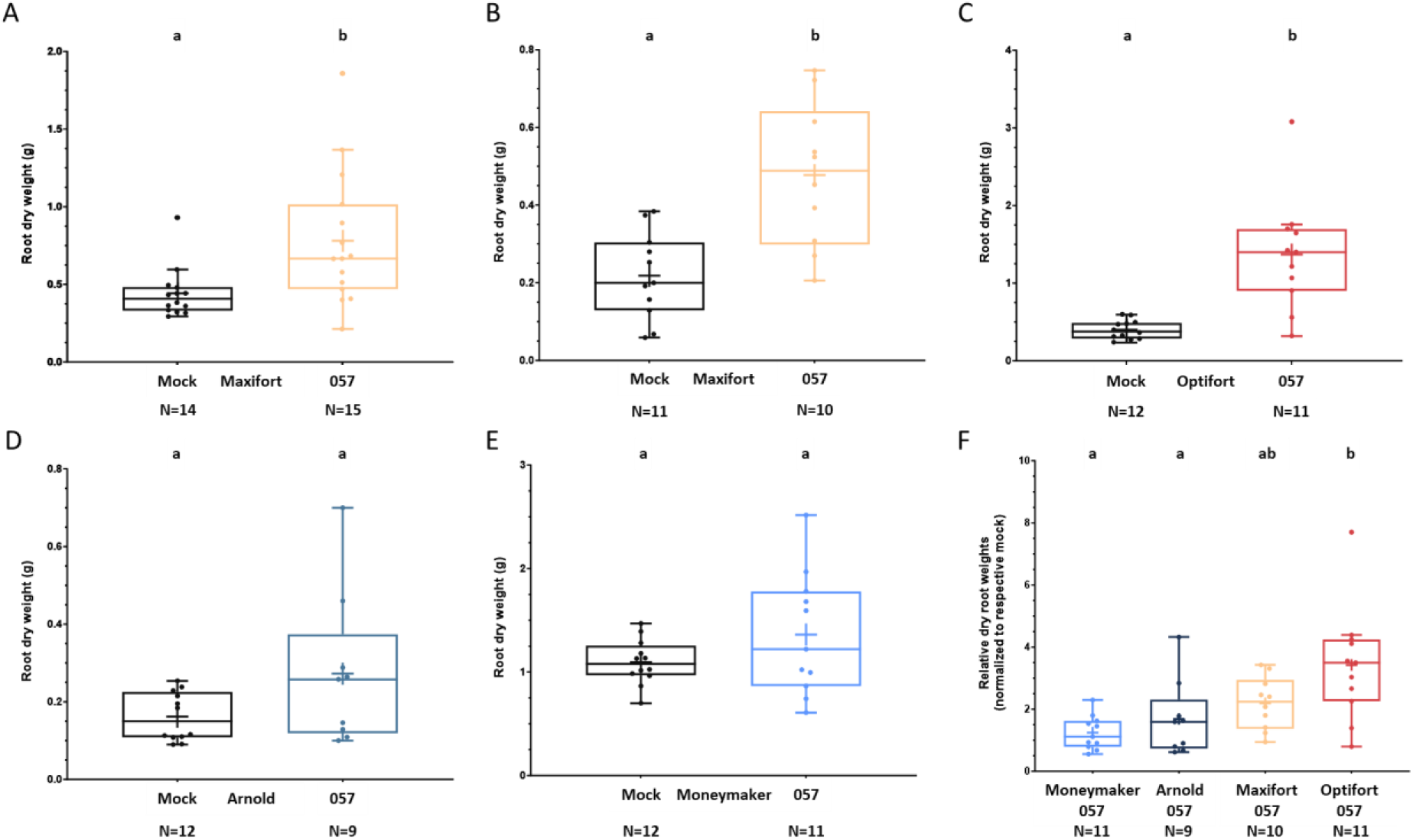
Comparisons of root dry weights of tomato and tomato rootstocks post inoculation with rhizogenic *Agrobacterium* strain 057 in a soil-based bioassay. Plants were harvested eight weeks post first inoculation, and root dry weights were measured. *A)* Pilot experiment using only ‘Maxifort’. Statistical analyses were performed using an unpaired t-test with Welch’s correction with two-tailed *p*-value. *B-E*) Main study to assess HRD in *B)* ‘Maxifort, *C)* ‘Optifort, *D)* ‘Arnold’, and *E)* ‘Moneymaker’. Statistical analyses were performed using unpaired t-test with Welch’s correction with two-tailed *p*-value. *F)* Relative root dry weights of plant varieties post inoculation with strain 057. The root dry weights of strain 057-treated plants were normalized to the mean values of their mock-treated counterparts. Statistical analyses were performed using parametric ordinary one-way ANOVA with Tukey’s multiple comparisons *post hoc* test. Different lowercase letters above bars indicate statistically significant differences between the groups.

We next conducted a larger study to compare the susceptibilities of ‘Moneymaker’, ‘Maxifort’, ‘Optifort’, and ‘Arnold’ against rhizogenic *Agrobacterium* strain 057. Plant roots were harvested eight weeks after the first inoculation, and root dry weights were recorded. Inoculation with strain 057 resulted in a significant 2.2-fold increase in root dry weight in ‘Maxifort’ and a 3.4-fold increase in ‘Optifort’ compared to their respective mock-treated controls (Fig. 2 B-C). In contrast, ‘Arnold’, and ‘Moneymaker’ showed no significant increase compared to their mock-treated controls (Fig. 2 D-E).

Because the employed genotypes differed substantially in average root dry weights, direct comparisons of the root dry weights between different genotypes were not appropriate. We therefore normalized root dry weights of strain 057-treated plants to the mean of their mock-treated counterparts. This analysis revealed that only ‘Optifort’ exhibited significantly greater susceptibility than both ‘Arnold’ and ‘Moneymaker’, showing 2-fold and 2.7-fold higher relative increases in root dry weights, respectively, while ‘Maxifort’ did not show significant differences compared to ‘Optifort’, ‘Arnold’, and ‘Moneymaker’ (Fig. 2F). These findings identified ‘Optifort’ as the most susceptible genotype among those assessed, followed by ‘Maxifort’, while ‘Arnold’ and ‘Moneymaker’ showed comparatively higher tolerance to rhizogenic *Agrobacterium* strain 057.

### 6.2 Comparative virulence testing of rhizogenic *Agrobacterium* strains

We next examined whether the soil-based bioassay could discriminate virulence levels among different rhizogenic *Agrobacterium* strains. To do so, we assessed the susceptibility of ‘Optifort’, the most susceptible genotype in our previous assay, to six rhizogenic *Agrobacterium* strains belonging to different genomospecies: 057 (genomospecies G25), K599 (genomospecies G21), 012 (genomospecies G9), 097 (genomospecies G9), and 4062 (genomospecies G7). These strains were originally isolated from HRD-affected plants in greenhouses (Bosmans et al. 2015), but their virulence levels have not been characterized in tomato plants and rootstocks. Additionally, a disarmed K599 derivative strain, 18r12v, which lacks the Ri T-DNA and does not induce HRD, was included as a negative control along with mock-treated controls. Furthermore, to capture potentially enhanced symptom development and/or differences among the tested strains, we extended the assay endpoint from nine to twelve weeks.

Inoculation with strains 057, K599 and 012 significantly increased root dry weights by 2.2-, 1.7-, and 1.4-fold, respectively, compared to the mock-treated controls (Fig. 3). In contrast, strains 4062, 097 and the disarmed strain, 18r12v, did not induce a significant increase in root dry weight (Fig. 3). Direct comparisons of relative root dry weights revealed that strain 057 induced significantly higher root dry weights than strains 097, 012, 4062, and 18r12v, while K599 induced significantly higher root dry weight than strains 097, 4062 and 18r12v (*SI Appendix*, Fig. S2). Strain 012 also yielded significantly higher root dry weight than strain 097 and 18r12v (*SI Appendix*, Fig. S2). These results demonstrate that ‘Optifort’ provides a suitable system for distinguishing differences in virulence among various rhizogenic *Agrobacterium* strains based on root dry weights.

**Fig. 3.**
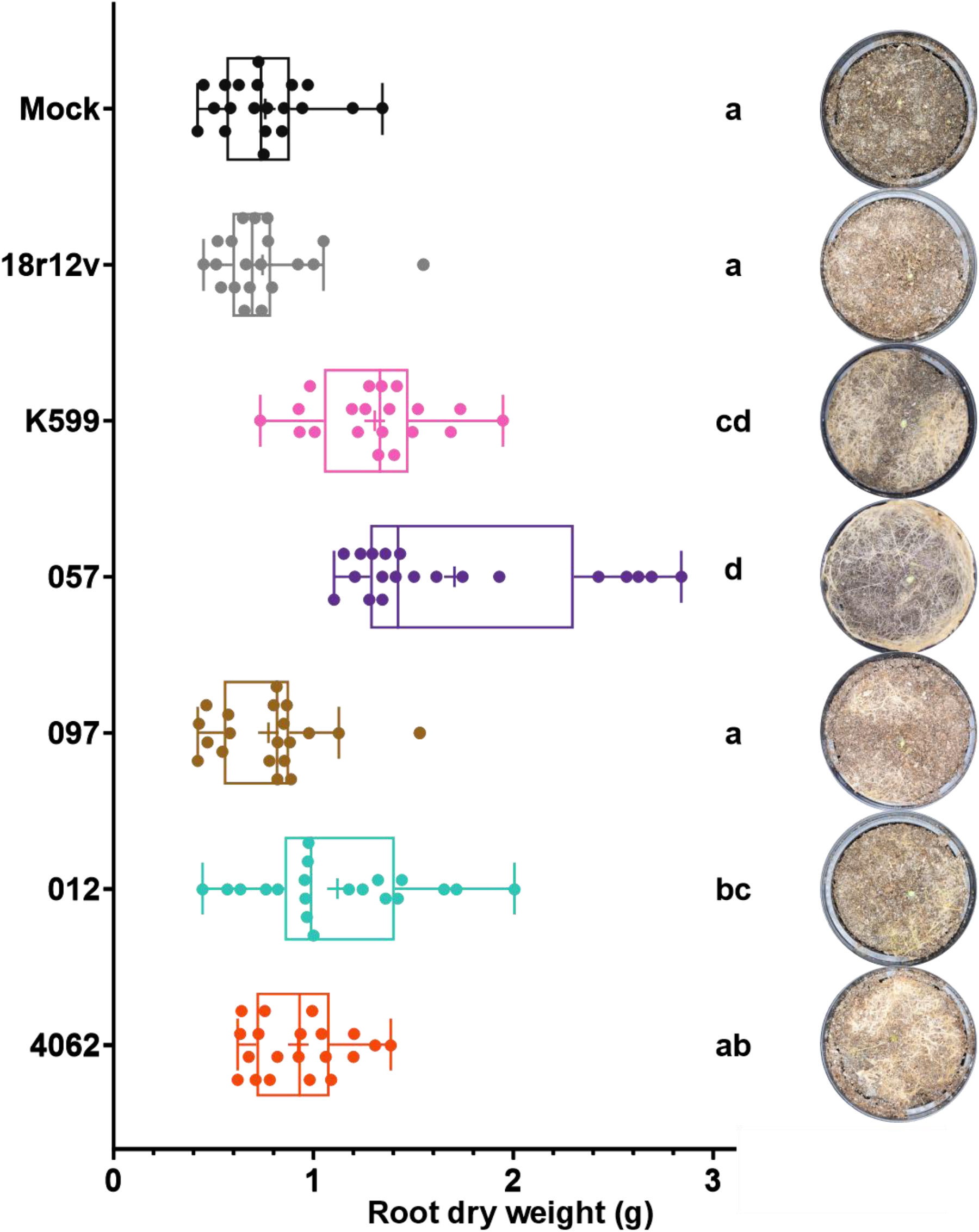
Comparisons of root dry weights and agravitropic root development for ‘Optifort’ plants inoculated with various rhizogenic *Agrobacterium* strains. Root dry weights and representative images of soil surface of ‘Optifort’ plants inoculated with rhizogenic *Agrobacterium* strains 18r12v, K599, 057, 097, 012 and 4062, eleven weeks post first inoculation. Statistical analyses were performed using parametric ordinary one-way ANOVA with Tukey’s multiple comparisons *post hoc* test. Different lowercase letters indicate statistically significant differences between the groups. N=20 for all treatments.

The employed strains also differed in their ability to induce root development near the soil surface i.e., agravitropic root growth (Fig. 3 and *SI Appendix*, Fig. S3). Inoculation with strains 057, K599, and 4062 showed pronounced root development at the soil surface compared to plants treated with strains 012, 097, 18r12v, or mock-treated controls. Notably, strain 4062 induced prominent agravitropic root development despite failing to induce a significant increase in root dry weight, whereas strain 012 showed increased root dry weight but produced fewer agravitropic roots. These contrasting outcomes suggest that agravitropic root development and root biomass represent partially independent HRD symptoms.

To confirm whether the agravitropically developing roots were genetically transformed, we analyzed the expression of the T-DNA gene *root oncogenic locus C* (*rolC*) which was found to be highly expressed in tomato hairy roots (Gryffroy et al. 2023), and normalized it to the reference gene Sl*CAC* (Expósito-Rodríguez et al. 2008). Sl*CAC* expression was consistent across treatments (Cq 22-25), while *rolC* expression varied widely (Cq ranges: K599: 17-28; 057: 19-20; 012: 17-32; 097: 34-35; 4062: 17-32; 18r12v and Mock: not detected). This variability likely reflects the presence of a composite root system in the infected plants comprising both, hairy and wild-type roots, which dilute T-DNA-related transcripts. Agarose gel electrophoresis of RT-qPCR amplicons confirmed *rolC* expression in all rhizogenic *Agrobacterium*-treated samples (*SI Appendix*, Fig. S4), indicating that even low virulent strains like 097 and 4062, generated transformed roots.

## 7 Discussion

HRD poses a serious constraint on the commercial production of Solanaceae (tomato, bell pepper, and eggplant) and Cucurbitaceae (cucumber and melon) crops globally (Bosmans et al. 2017b). HRD primarily infects the root system of the cultivated plant, particularly the rootstock in grafted plants. However, only limited and mostly qualitative information is available on HRD susceptibility of tomato cultivars and rootstocks under natural infection settings (de Freitas and Taylor 2023; Ludeking and Janse 2011; Vanlommel et al. 2019). Our results show that a soil-based, non-wounding disease bioassay reliably reproduces HRD symptoms under ecologically relevant conditions and supports quantitative assessments of disease symptoms. Our findings are consistent with previous qualitative observations in tomato rootstocks (Ludeking and Janse 2011; Vanlommel et al. 2019) and highlight the value of an ecologically relevant assay for studying HRD under controlled conditions.

Root dry weight was found to be a consistent and informative quantitative marker for investigating tomato HRD. In response to rhizogenic *Agrobacterium* treatment, ‘Optifort’ and ‘Maxifort’ exhibited greater increases in root biomass compared to ‘Arnold’ and ‘Moneymaker’, confirming that tomato cultivars and rootstock varieties differ significantly in HRD symptom development. These differences are in agreement with earlier reports describing variable symptom development across different rootstocks (Ludeking and Janse 2011; Vanlommel et al. 2019). Since the tested rootstock genotypes share *S. lycopersicum* x *S. habrochaites* genetic backgrounds, our results suggest that specific parental lines strongly influence HRD susceptibility. Similar genotype-specific variability in tomato rootstocks has also been reported for other stresses, including *Verticillium* wilt, root knot nematodes, or salinity (Cortada et al. 2008; Gioia et al. 2013; Papadaki et al. 2017). Identifying genetic factors contributing to these differences may support the development of future rootstocks that balance vigor, (a)biotic stress tolerance, fruit yield, and reduced HRD susceptibility. Within this framework, ‘Optifort’ may serve as a susceptible genotype for future screening efforts, while ‘Arnold’ may be employed as a tolerant variety.

The bioassay also enabled comparative assessment of virulence among different rhizogenic *Agrobacterium* strains. Strains 057, K599, and 012 induced significantly higher root biomass compared to strains 4062 and 097. Extending the assay from nine to twelve weeks further revealed pronounced agravitropic root development for some strains. Interestingly, the extent of agravitropic root development did not correlate consistently with increased root biomass. For example, strain 4062 showed prominent agravitropic root development without a significant increase in root biomass, while strain 012 induced fewer agravitropic roots but showed a significant increase in root biomass. These observations suggest that HRD manifests along at least two partially independent axes: *a)* increased root biomass and *b)* agravitropic root development. Future work should determine which HRD symptoms most strongly impact crop performance, since growers may be hindered by *a)* increased root biomass that diverts resources away from fruit production; or *b)* extensive root proliferation that interferes with irrigation (Bosmans et al. 2017b; de Freitas and Taylor 2023).

Expression analyses of the T-DNA gene *rolC* confirmed that all tested rhizogenic *Agrobacterium* strains, including those with low virulence, successfully transferred T-DNA to the host plant. We analyzed *rolC* expression because *a) rolC* is only expressed in plants, and *b) rolC* was found to be highly expressed in our previous transcriptome studies on tomato hairy roots (Gryffroy et al. 2023). Variation in *rolC* transcript abundance in the harvested roots likely reflects composite nature of the infected root system, which includes both transformed hairy roots, and non-transformed wild-type roots. These findings show that strains inducing negligible or mild root biomass increases are nonetheless competent at T-DNA transfer and may induce subtle HRD symptoms.

This assay has practical applications for future resistance breeding and HRD management. It enables systematic screening of rootstocks or parental lines to identify susceptible or resistant genotypes. The assay is also suitable for evaluating biocontrol strategies, including bacteriophages, *Pseudomonas, Paenibacillus*, and *Rhodococcus* strains, that reduce rhizogenic *Agrobacterium* populations or reduce HRD symptom development (Bosmans et al. 2017a; Bourigault et al. 2021; Fortuna et al. 2023; de Freitas and Taylor 2023; Vargas et al. 2021). Additional approaches such as essential oils or the cyclic lipopeptide plipastatin, previously shown to reduce crown gall symptoms, may also be explored (Brown et al. 2025; Habbadi et al. 2018).

Future studies can integrate image-based quantification of agravitropic roots as demonstrated for HRD-affected bell peppers (Eberle et al. 2020) to reduce manual labor. Additionally, reporter genes such as GFP, mCherry or RUBY might also be included to differentiate and quantify Ri T-DNA transformed roots from non-transformed roots (de Freitas and Taylor 2023; Rodrigues et al. 2025). Such tools could further refine HRD phenotyping and possibly uncover additional parameters linked to HRD. As whole genome sequencing of rhizogenic *Agrobacterium* strains expands (Kim et al. 2023; Otten 2021; Vargas et al. 2024; Weisberg et al. 2020), pairing genomic comparisons to phenotypes derived from this assay may help identify virulence determinants influencing HRD severity. Such insights may also guide plant transformation technologies, given the utility of highly virulent strains like K599 in species such as soybean as well as regenerative potential of hairy roots for obtaining transformed plants (Duan et al. 2025; Goralogia et al. 2025).

Overall, our soil-based assay provides an ecologically relevant framework for studying tomato HRD, comparing virulence among rhizogenic *Agrobacterium* strains, and supporting the development of effective management strategies in hydroponic tomato production systems.

## 3. Abbreviations list

ANOVA: Analysis of Variance
cDNA: Complementary deoxyribonucleic acid
CFU: Colony forming units
Cq: Quantitative cycle
cv.: Cultivar
HRD: Hairy root disease
MES: 2-(N-morpholino)ethanesulfonic acid
MS: Murashige and Skoog medium
ORF: Open reading frame
RT-qPCR: Reverse transcriptase quantitative polymerase chain reaction
Ri: Root-inducing
*rolC*: *Root oncogenic locus C* gene
Sl*CAC*: *Solanum lycopersicum CLATHRIN ADAPTOR COMPLEXES MEDIUM SUBUNIT* gene
T-DNA: Transfer DNA
V/V: Volume/volume
WGS: Whole genome sequencing
YEP: Yeast extract peptone medium

## 8. Acknowledgements

We would like to thank Lut Ooms (Division of Crop Biotechnics, Biosystems Department, KU Leuven), ir. Poi Verwilt and Loïck Derette (Greenhouse Division, Biosystems Department, KU Leuven) for providing technical advice and assistance for this research and ir. Anna Bens and ir. Ward Boven (Division of Crop Biotechnics, Biosystems Department, KU Leuven) for critically reading the manuscript. We would also like to thank dr. Roger Thilmony and the United States Department of Agriculture-Agricultural Research Service (USDA-ARS) for providing strain 18r12v-Δ*recA*::*tetA*.

## 9. Funding

S.D.R. is supported by KU Leuven Postdoctoral Mandate (PDMT2-22-035) and Research Foundation – Flanders Junior Postdoctoral Fellowship (1S43918N, 1S43920N, 12AKQ24N). The work was also supported by KU Leuven internal funding (C14/19/074).

## 10. Ethics Declarations

### Ethics approval and consent to participate

All experimental research conducted in this study complied with relevant institutional, national, and international guidelines and legislations.

### Consent for publication

Not applicable.

### Competing interests

The authors of this study declare no competing interests.

## 11. Author information

### Author contributions

S.D.R., L.P., and B.D.C. designed experiments; N.K. and H.R. provided bacterial strains used in the study and contributed to discussions; S.D.R., N.K., and J.M. performed experiments; S.D.R. and B.D.C. analyzed the data; S.D.R., H.R., L.P., and B.D.C. wrote the manuscript. All authors revised the manuscript.

## 12. Availability of Data and Materials

Not Applicable

## Notes

### Competing Interest Statement

The authors have declared no competing interest.

