## Supplementary Information Appendix for "Soil-based disease bioassay for the study of rhizogenic *Agrobacterium*-tomato interactions"

| | |
|---|---|
| Savio Rodrigues | 0000-0003-1641-6310 |
| Nuri Kim | 0000-0002-5084-6448 |
| Joram Moons | |
| Laurens Pauwels | 0000-0002-0221-9052 |
| Hans Rediers | 0000-0001-8645-107X |
| Barbara De Coninck | 0000-0002-9349-5086 |

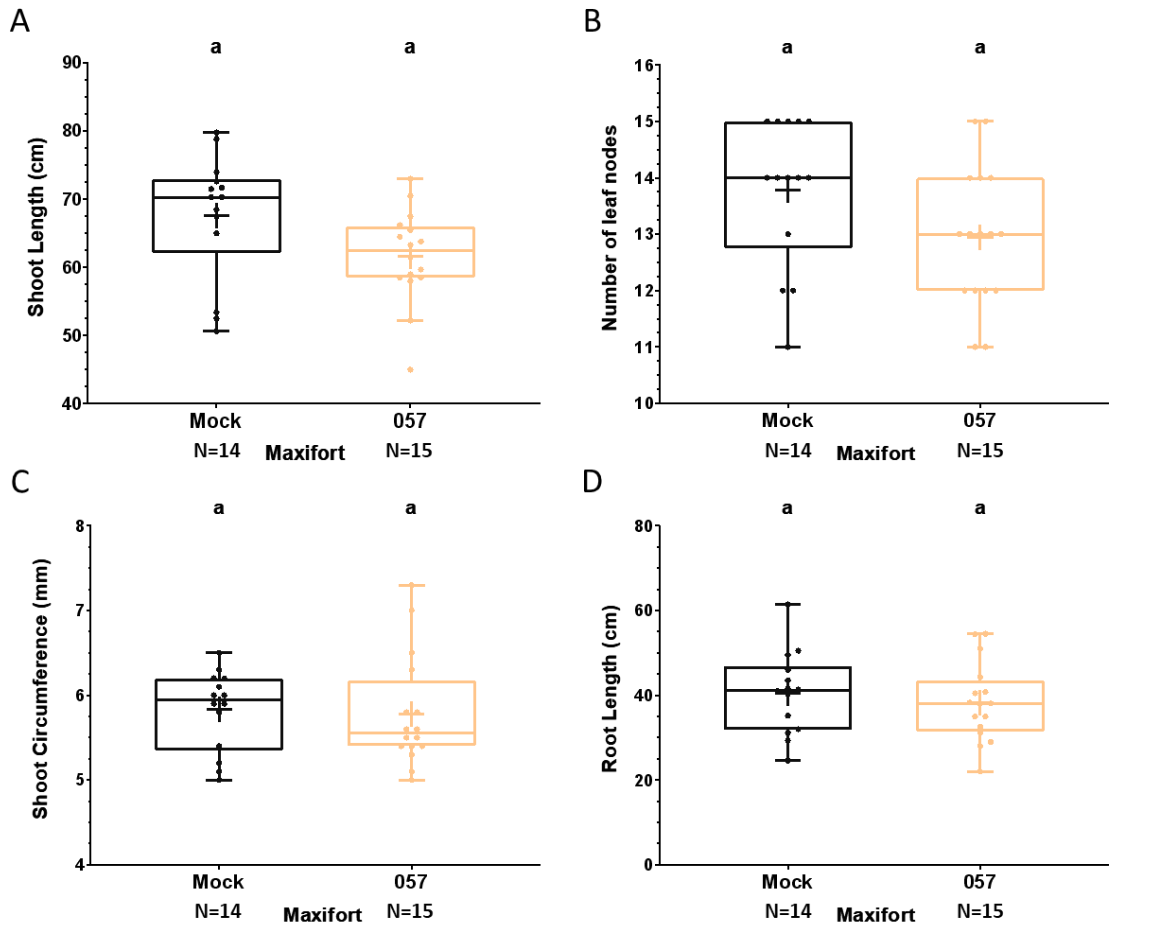


**Fig. S1: Measurement of phenotypic parameters of 'Maxifort' (*Solanum lycopersicum* x *Solanum habrochaites*) plants 8 weeks post inoculation with rhizogenic *Agrobacterium* strain 057.** *A)* Comparison of shoot lengths of mock- and 057-inoculated plants in centimeters (cm). *B)* Comparison of number of leaf nodes of mock- and 057-inoculated plants in centimeters (cm). *C)* Comparison of shoot circumference of mock- and 057-inoculated plants in millimeters (mm). *D)* Comparison of shoot lengths of mock- and 057-inoculated plants in centimeters (cm). Statistical analyses were performed using unpaired t-test with Welch's correction and two-tailed *p*-value.

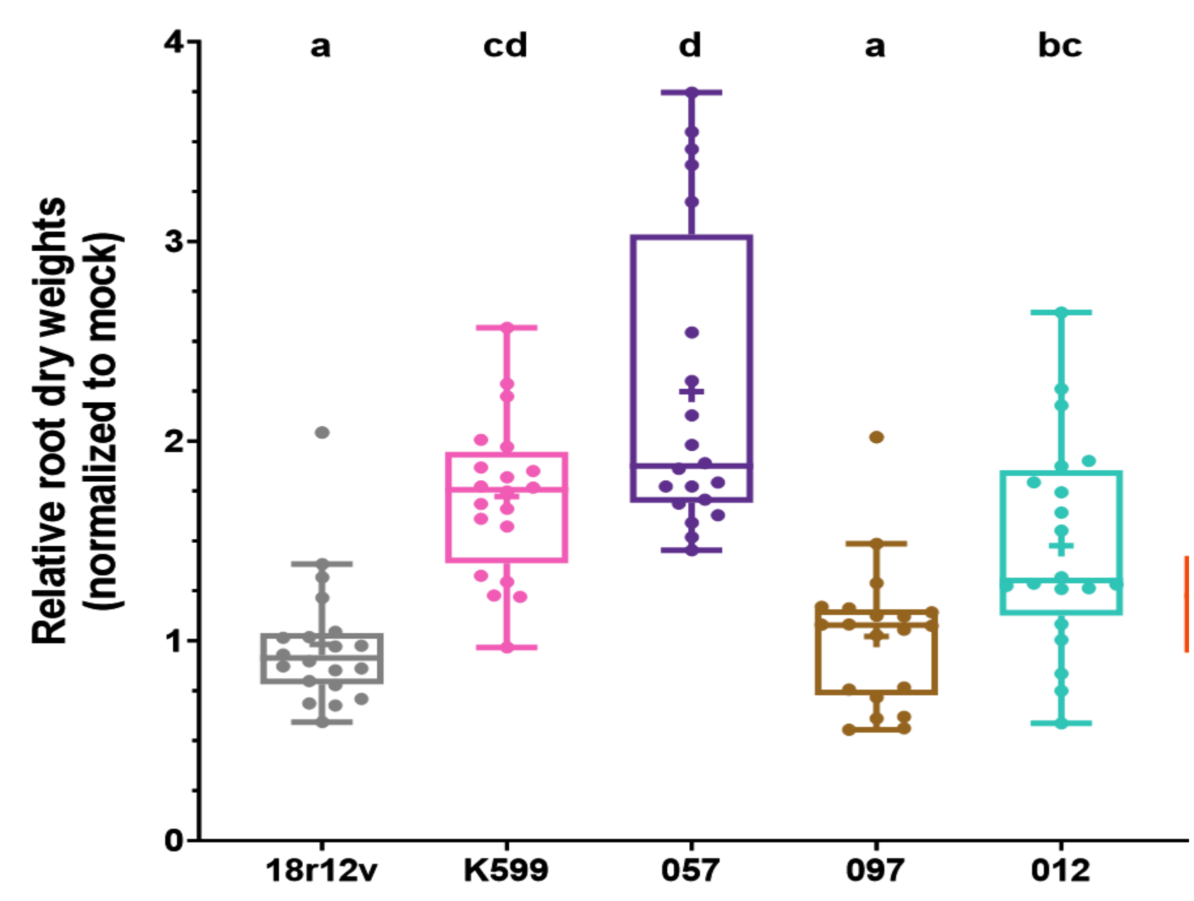


**Fig. S2: Comparisons of relative fold-changes in root dry weights of 'Optifort' (*S. lycopersicum* x *S. habrochaites*) plants inoculated with various rhizogenic *Agrobacterium* strains.** Relative root dry weights of 'Optifort' plants inoculated with rhizogenic *Agrobacterium* strains 18r12v, K599, 057, 097, 012 and 4062, eleven weeks post first inoculation. The root dry weights of rhizogenic *Agrobacterium*-treated plants were normalized to the mock. Statistical analyses were performed using parametric ordinary one-way ANOVA with Tukey's multiple comparisons post hoc test. Statistical significances are denoted by alphabetical letters 'a', 'b', 'c' and 'd'. N=20 for all treatments.

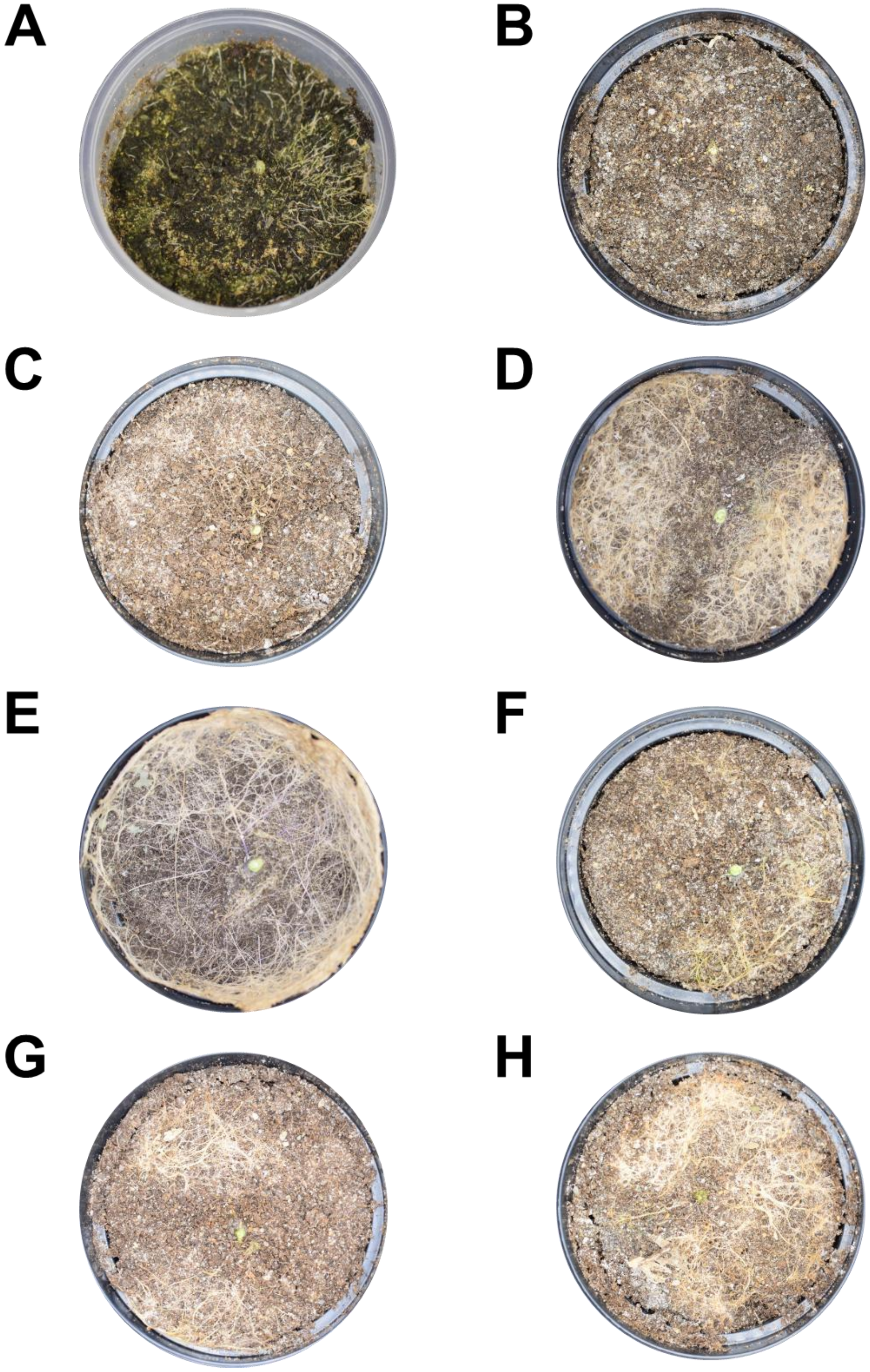


**Fig. S3: Comparisons of agravitropic root development in 'Optifort' inoculated with various rhizogenic Agrobacterium strains.** *A)* Representative image of soil surface of 'Optifort' plant inoculated with strain 057 eight weeks post first inoculation. *B-H)* Representative images of soil surface of 'Optifort' plants inoculated with *B)* Mock, *C)* 18r12v, *D)* K599, *E)* 057, *F)* 012, *G)* 097 and *H)* 4062 eleven weeks post first inoculation.

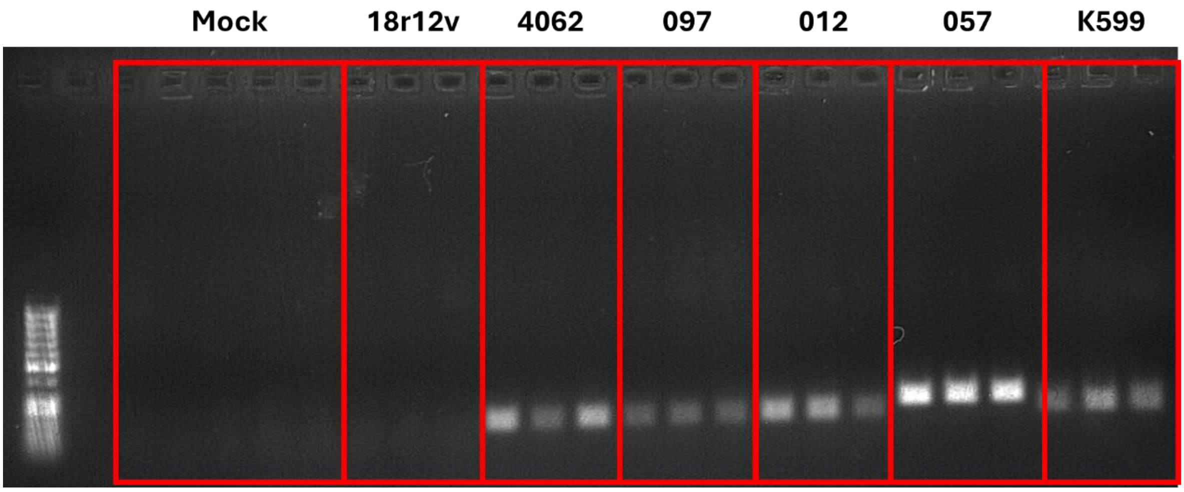


**Fig. S4. Expression of *rolC* in 'Optifort' treated with rhizogenic *Agrobacterium*.** Quantitative reverse-transcriptase polymerase chain reaction (RT-qPCR) was performed to assess the expression of *rolC* in RNA samples obtained from the roots of plants which were subject to inoculations with 1) Mock, 2) 18r12v, 3) 4062, 4) 097, 5) 012, 6) 057, and 7) K599. After RT-qPCR, the samples were loaded on an agarose gel to visualize the *rolC* amplicon.
